# Intrinsic Disorder and Degeneracy in Molecular Scale Organization of Biological Membrane

**DOI:** 10.1101/582338

**Authors:** Sahithya S. Iyer, Anand Srivastava

## Abstract

The scale-rich spatiotemporal organization in biological membrane dictates the “molecular encounter” and in turn the larger scale biological processes such as molecular transport, trafficking and cellular signalling. In this work, we explore the degeneracy in lateral organization in lipid bilayer systems from the perspective of energy landscape theory. Our analysis on molecular trajectories show that bilayers with lipids having *in-vivo* characteristics have a highly frustrated energy landscape as opposed to a funnel-like energy landscape in *in-vitro* systems. Lattice evolution simulations, with Hamiltonian trained from atomistic trajectories using lipids topology and non-affine displacement measures to characterize the extent of order-disorder in the bilayer, show that the inherent frustration in *in-vivo* like systems renders them with the ability to access a wide range of nanoscale patterns with equivalent energy penalty. We posit that this structural degeneracy could provide for a larger repository to functionally important molecular organization in *in-vivo* settings.

---

From the emerging field of lipidomics research, it is now known that there are more that 37,500 unique lipid species in Eukaryotic and Prokaryotic cells. The plasma membrane itself has more than 500 different molecular species of lipids.^1–5^ One of the fundamental questions in the field is “Why are there so many lipids”?^6,7^ From being considered simply as a “selectively permeable barrier” made up of lipid bilayer,^8–10^ it is now well established that lipids in biological membrane have a wide range of functional roles.^11–15^ Lipid environments modulate innumerable biological processes^16–27^ and in certain cases, specific lipids are critical for a given biological function. For example, protein p24 in COPI machinery recognizes a single sphingolipid species to induce oligomerization and vesicle transport in the ER.^28^ Another example in which lipids play a crucial role is that of the very rare highly phosphatidylinositol lipid which drive activation of K+ channels^29,30^ besides several other biological functions by specific interactions.^31,32^ In all the examples discussed above, lipid interactions range from being extremely non-selective to highly discriminating^33–35^ and such a range of interaction is possibly facilitated by their large diversity and chemical complexity.

Though the rationale behind diversity in lipid structures in biological membranes is appreciated from the examples related to the folding, dynamics and functions of membrane proteins^36–40^ and other molecular-scale processes such as drug binding,^41^ and lipid defects driven selective peptide partitioning,^42,43^ the evolutionary advantage of such a huge variety and complexity in lipid species is yet to be fully understood especially in the light of the high metabolic expense of lipid synthesis and the cost of maintaining the required lipid homeostasis in the cellular membranes. With the advent of super-resolution imaging techniques, there is now compelling evidence proving that lipid mixtures, like simple binary alloy systems (AB) show non-random demixing and phase separation in both *in-vitro* and *in-vivo* scenarios.^44–48^ However, unlike the simple binary alloy mixtures that show phase co-existence consisting of isolated ‘A rich’ and ‘B rich’ regions at thermal equilibrium, lipid mixtures show a plethora of interesting lateral organization, which can be attributed to the complex molecular interactions arising from differences in chemical structures of the lipid species.^49–54^ The sub-100 nm transient *patterns* on the biological membrane, which are generally stabilized far away from equilibrium in cells, are believed to be functionally important in various physiological processes.^55–59^ Signal transduction^60^ localization of biochemical processes,^61^ endocytosis and exocytosis of membrane proteins,^62^ organelle marker for specific proteins^63,64^ and phases of cell cycle^65^ are some examples of such processes.

Given the functional importance of the lateral organization in biological membranes, an immediate question that arises in the context of a complex lipidome is: Can the lateral organizations formed on membrane surface be degenerate in certain conditions? Degeneracy in this context refers to the ability of the lipids to organize into different “pattern” on the membrane surface with minimal energy penalty. ^66^ We ask this question because degeneracy in membrane lateral organization could be a possible consequence of the complex and abundant lipidome. We posit that similar to recognition motifs on protein surfaces involved in functions ranging from sensing lipid packing and curvature in the membrane^67^ to specific binding to DNA,^68^ the unique lateral organization patterns formed by combination of lipid species might function as platform for a range of physiologically important processes such as protein partitioning, intracellular membrane trafficking and “parallel processing” of signalling events.^69^ The degeneracy in membrane lateral structure could be an evolutionary design principle that maximizes functional output with a given set of lipid species.

In this work, we focus on degeneracy in lateral organizations using three model ternary lipid systems: DPPC/DOPC/CHOL, PSM/DOPC/CHOL and PSM/POPC/CHOL, respectively. The composition in each system is carefully chosen such that the system has coexisting liquid order (*L_o_*) and liquid disorder (*L_d_*) regions.^70,71^ While the first two systems are more commonly studied in *in-vitro* set-ups, the last system has a lipid types closer to that in *in-vivo* environment. Considering the functional importance of variety in lateral organizations, we explore if the given ternary mixture of lipids can adopt different organizations with similar energy penalty. Towards this, we first post-process some very long atomistic simulation trajectories of the three bilayer systems with fluid phase co-existence.^72,73^ The systems information is provided as Table 1. We then calculate the interaction energies between lipids as a function of a newly defined order parameter (*χ*^2^) that utilizes the information on topological changes and displacements of neighbouring lipids to locally characterize the extent of order/disorder in the membrane without any knowledge of the chemical identity of the lipids.^74,75^ Formulations and schematic related to measurement of non-affine (*χ*^2^) value (Figure S1a) and details of calculations of interaction energy (Figure S1b) is discussed in the supporting information (SI). The *χ*^2^ values of the three phase separated systems shows a bimodal distribution and is able to capture the ordered and disordered phases without the knowledge of chemical identity of the lipids (Figure S2 and S3). We also provide comparisons of (*χ*^2^) against measures such as deuterium order parameter and hexatic order parameter to highlight the advantages of characterizing molecular-level phase coexistence in lipid bilayer systems using χ^2^ measures (Figure S4a and S4b). This characterization was first used as a measure of shear stress propagation in metallic glasses to study irreversible plastic deformations ^76^ and have been extensively applied recently on soft matter systems^77–79^ and on biological systems.^74,75,80,81^ Figure S5 shows the calculated interaction energies for each lipid as a function of its *χ*^2^ values for the three systems of interest. We observe that degree of positive correlation between lipid-lipid interaction energies and *χ*^2^ values gradually decreases from DPPC/DOPC/CHOL to PSM/DOPC/CHOL and PSM/POPC/CHOL. This has a direct influence on the extent of phase separation resulting in separate clusters of different morphologies of *L_o_* and *L_d_* species in the three systems.

The profiles shown in Figure 1 can be treated as an analogue of potential energy surface (PES) with *χ*^2^ as its reaction coordinate. A better graphical visualization of the PES is through an energy (microcanonical) disconnectivity graph.^82–85^ Each vertex in the graph represents an energy minimum that we call basin and all the basins that are connected by barriers less than *k_B_T* (a choice of Δ*E* appropriate for physical systems) represent a super basin. The super basins are disjoint open sets and hence the name disconnectivity graph. Figure 1 shows the energy disconnectivity graphs for the three systems. Moving along the direction of the graph edges (downwards), DPPC/DOPC/CHOL and PSM/DOPC/CHOL systems have a distinct funnel like overall shape, while PSM/POPC/CHOL has multiple minima with similar energies. These equivalent minima are separated by high energy barriers indicating frustration in this system.^86^ Hence, presence of multiple minima of similar energies allows for a higher variability in lateral organization patterns with similar energy penalty. We propose that propensity of PSM/POPC/CHOL system to have a wide array of minimum energy states favours selection of these lipids in natural system over others in the huge milieu of possible lipid structures. In short, we find that the interaction energy landscapes in ternary lipid systems ranges from funnel like for systems with *in-vitro* like compositions to frustrated for systems with *in-vivo* like compositions.

**Figure 1:**
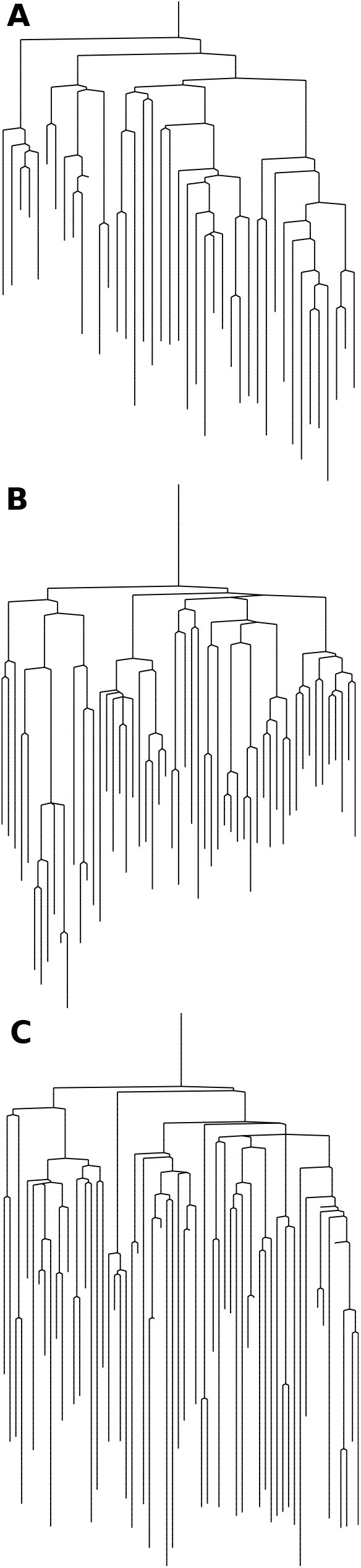
Energy disconnectivity graph of (A) DPPC/DOPC/CHOL, (B) PSM/DOPC/CHOL and (C) PSM/POPC/CHOL systems with energy spacing of 1 *k_B_T*. Vertical axis denotes energy and horizontal axis *χ*^2^ values.

To test preliminary hypothesis regarding patterns formed in lateral organizations in the three ternary lipid mixtures, we evolve a lattice model with a Hamiltonian that is trained from the all-atom (AA) trajectories data. The ternary lipid systems are mapped to a lattice model where each lattice site is assigned a *χ*^2^ value from a chosen distribution (figure S3). Interaction energy between a pair of nearest neighbour sites is written as

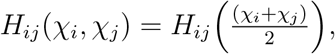

where *i,j* are the first four nearest neighbours on the lattice configuration. The interaction energy profiles are made continuous by performing a cubic interpolation across energy values for sites within a cut-off of 14Å. Interaction energies for lipid pairs separated by a cut-off of greater than 14Åis set to a higher disallowed energy value. The profiles thus generated for pairs of lipids are shown in Figure 2. We use enthalpic interaction energies to dictate lateral organization in the lattice model for the three ternary lipid systems with the assumption that phase separation in lipid mixtures at a given temperature are predominantly enthalpy driven. Details of lattice evolution performed using simulated annealing Monte Carlo algorithm in the provided in the SI. While the equilibrium organization in AA system is fairly well recapitulated in lattice models of DPPC/DOPC/CHOL (Figure 3) and PSM/DOPC/CHOL (Figure S6) systems, the lattice model for PSM/POPC/CHOL permits a broader range of spatial patterning (Figure 4). Figure 3 shows clumping behaviour of low *χ*^2^ values in DPPC/DOPC/CHOL system forming a single *L_o_* cluster, irrespective of the starting configuration and final configuration from lattice evolution is similar to the organization observed in equilibrium AA system. Different morphologies observed in PSM/POPC/CHOL system is a manifestation of its highly rugged average interaction energy profile. This is because the interaction energy profile does not preclude lipids with low *χ*^2^ values to coexist with lipids having higher *χ*^2^ values, thus resulting in a heterogeneous lateral phase organization. Due to the highly rugged nature of inter-lattice interactions, the final lattice configuration is path-dependent and hence depends on initial conditions. The final organization of *χ*^2^ values obtained from different initial starting configurations have similar energies as shown in figure 4. This indicates degeneracy in lateral organization patterns in this system. Experimental results from super resolution technique^46,47,87,88^ and single particle tracking diffusion measurements^89^ have shown that smaller domains with highly complex morphologies, resulting from non-ideal mixing, are observed in both lipids only systems and lipids-proteins bilayer systems. Our analysis clearly shows the diversity in lateral organizations for physiologically more relevant PSM/POPC/CHOL ternary system. We observe decreasing repulsive interactions between lipids with *L_o_* and *L_d_* signatures in the following order: DPPC/DOPC/CHOL > PSM/DOPC/CHOL > PSM/POPC/CHOL, which is consistent with previously noted in FRET experimental studies.^90–92^

**Figure 2:**
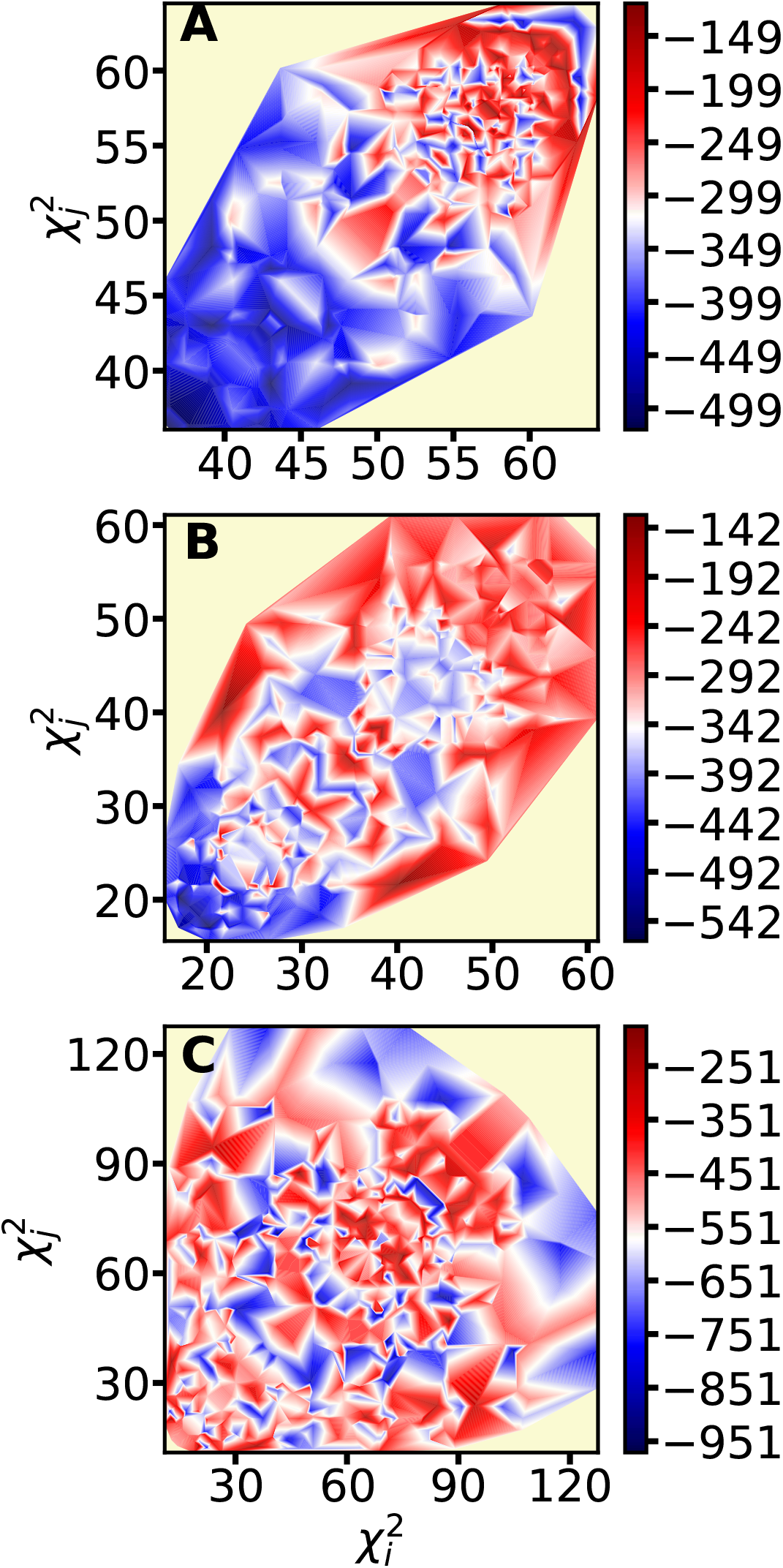
Average enthalpy of interaction written as 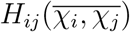 for (A) DPPC/DOPC/CHOL, (B) PSM/DOPC/CHOL and (C) PSM/POPC/CHOL systems. The pale yellow colors indicate disallowed energy regions during the MC sampling.

**Figure 3:**
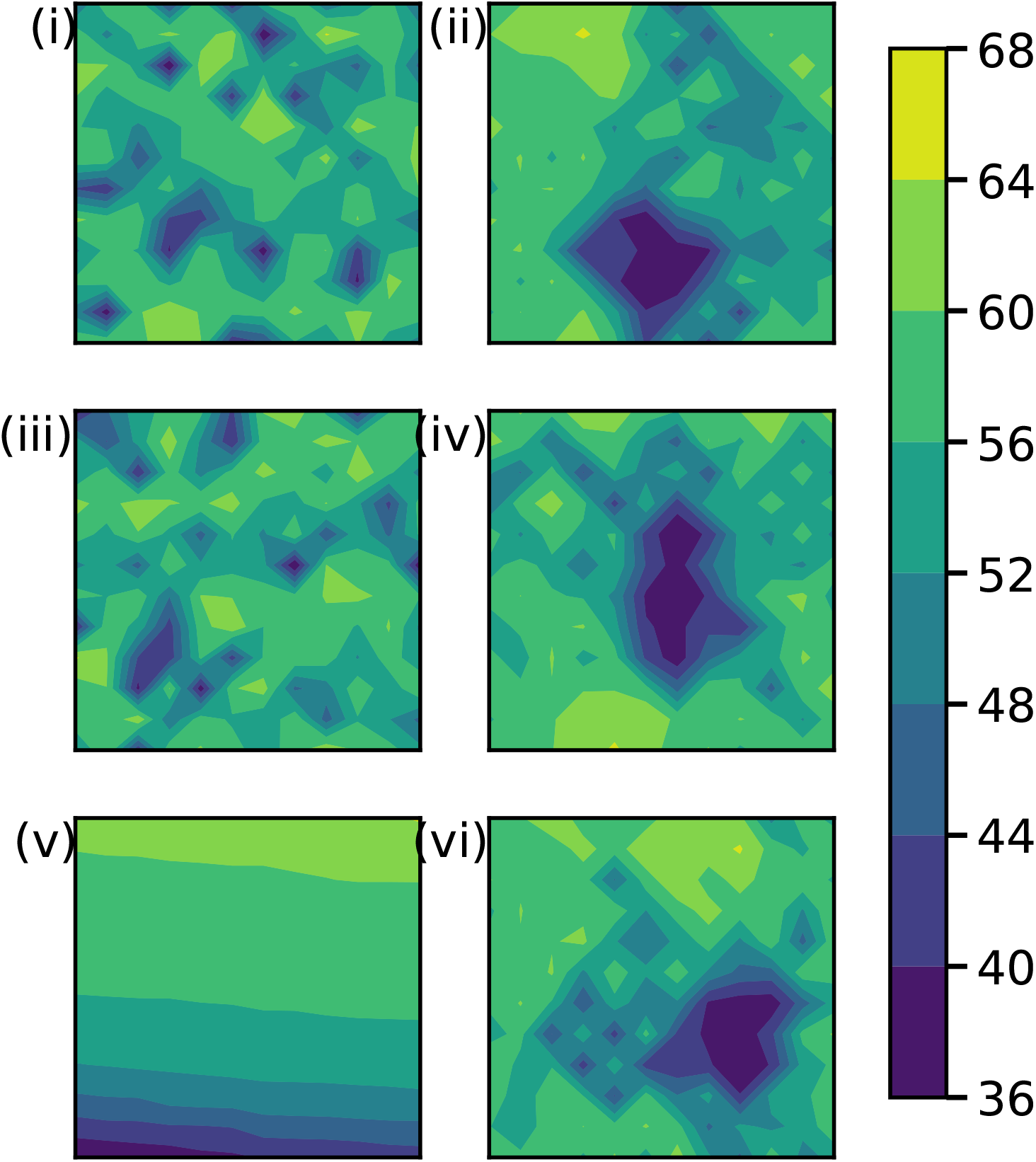
Initial (i,iii,v) *χ*^2^ organization and final (ii,iv,vi) *χ*^2^ organization for DPPC/DOPC/CHOL mixture. (i)-(ii) and (iii)-(iv) represent lattice evolutions starting from random lattice arrangements. (v)-(vi) represent lattice evolution from a phase separated arrangement of *χ*^2^ values. The sites have been color coded according to the *χ*^2^ values.

**Figure 4:**
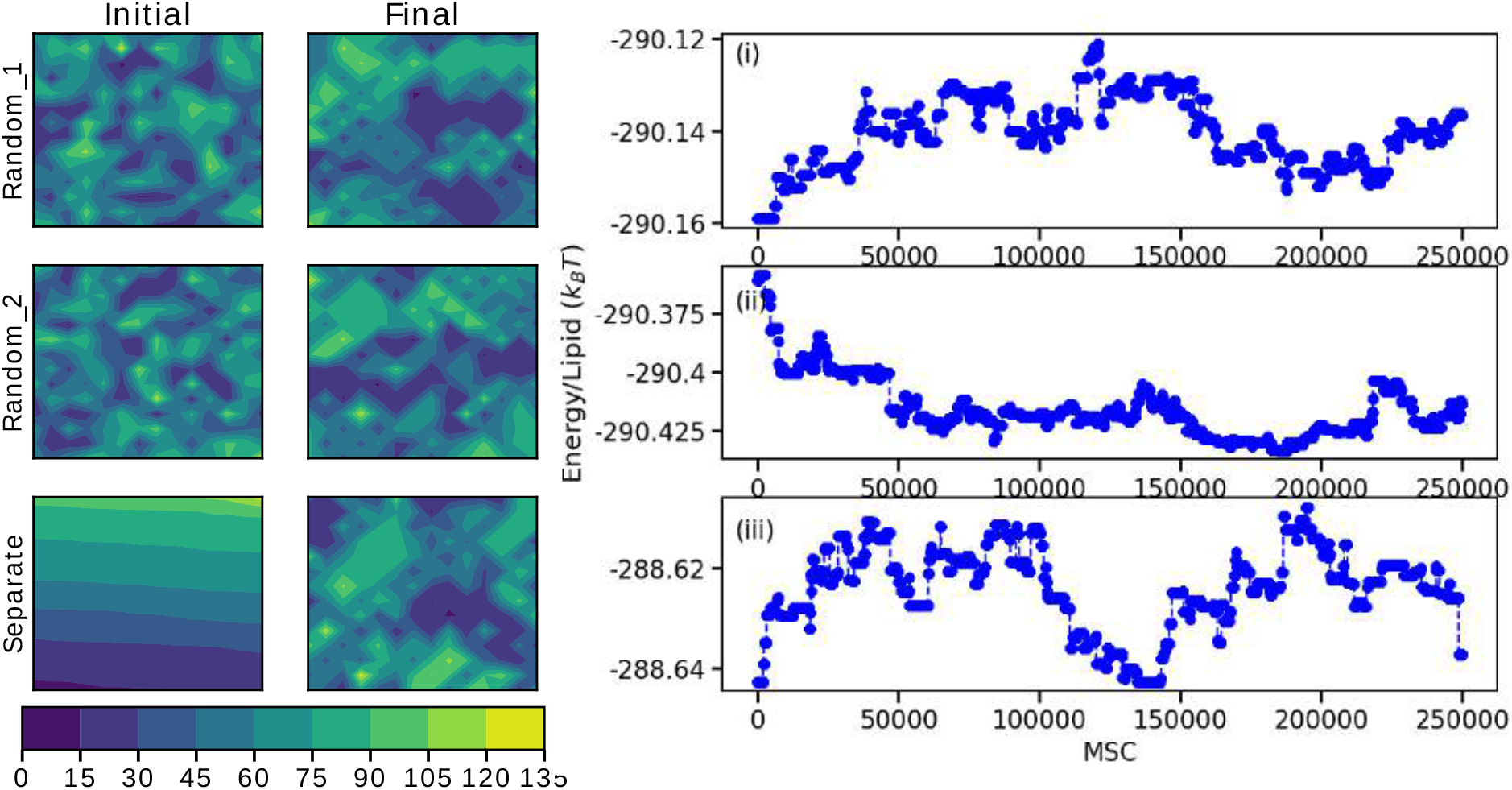
Initial *χ*^2^ organization and final *χ*^2^ organization for PSM/POPC/CHOL mixture. Row 1 and row 2 represent lattice evolutions starting from random lattice arrangements. Row 3 represent lattice evolution from a phase separated arrangement of *χ*^2^ values. The sites have been color coded according to the *χ*^2^ values. The Energy/lipid of last 2.5 * 10^5^ MCS evolution corresponding to evolutions shown in row (1,2 and 3) are shown in (i), (ii) and (iii) respectively.

We wanted to look into the sensitivity towards variety in lateral organization patterns as a function of lipid composition. Here we speculate that that are physiologically more relevant system would have access to a higher repertoire of molecular-scale lateral organizations as a function of their composition. Towards this, the ratio of saturated and unsaturated lipids is varied in the lipid mixtures as shown in figure S7 and the geometries of resultant clusters upon lattice evolution are recorded. Figure S8 shows that this ability to shuttle between different patterns of lateral organizations is clearly absent in DPPC/DOPC/CHOL system. Increasing the ratio of saturated lipids cause an increase in cluster size of the *L_o_* patch, but does not result in change in overall domain morphology. However, PSM/POPC/CHOL possesses a high degree of plasticity in its organization as evident from Figure 5. The changed ratios of *L_o_:L_d_* lipids for this system is shown in Figure S9.

**Figure 5:**
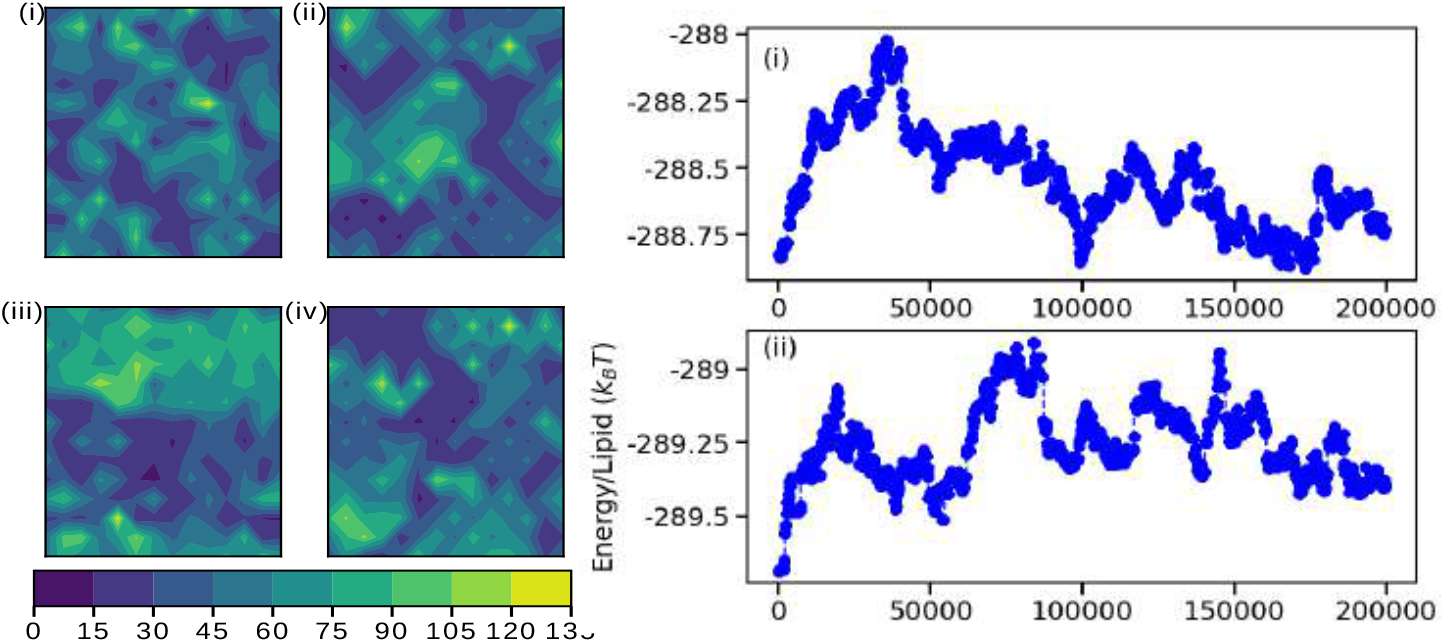
Initial (i,iii) *χ*^2^ organization and final (ii,iv) *χ*^2^ organization for PSM/POPC/CHOL mixture. (i-ii) represent lattice evolutions starting from random lattice arrangements and (iii-iv) evolution from phase separated lattice arrangement for *L_o_*:*L_o_* ratio of 0.72:0.28. The sites have been color coded according to the *χ*^2^ values. The Energy/lipid of last 2.0 * 10^5^ MCS evolution corresponding to evolutions is shown in the right.

Segregation and organization patterns on the lateral surface of cell membrane of living cells is primarily controlled through two mechanisms: (i) complex inter-lipid, inter-protein and lipid-protein interactions amongst chemically different lipid and protein species, and (ii) non-equilibrium membrane fluctuations mediated by underlying actin-myosin cortical region. In this work we have focused on lateral heterogeneities arising in pure lipid systems driven by inter-lipid interactions. The lipids in PSM/POPC/CHOL bilayer, which are physiologically relevant, possess the unique ability to access several degenerate organization patterns. In the context of evolutionary design principles, this seems highly relevant since biological membrane are optimally designed to minimize the cost of lipid synthesis and homeostasis and maximize the functions. Though there is a lack of compelling experimental evidence where signalling molecules are shown to directly sense domain morphology at molecular scales, there are examples of proteins that are known to modulate membrane organization and shape signal transduction across membrane and functionally depend on local membrane “fingerprints”.^93?,94^

Reverse scenario of protein preferentially partitioning into either *L_o_* or *L_d_* domain or at the *L_o_-L_d_* junction of a phase coexisting system is also observed.^95,96^ Stretching this observation further, we suggest that each of the cluster geometries in the lateral organization patterns can be considered to be a “recognition motif” or “signalling platform” similar to special stretches of amino acid sequences in proteins. The ability of physiologically relevant lipid composition mixtures to shuttle between a large number of lateral organization patterns enlarges the possible “query” space of recognition motifs. Though speculative, we believe the hypothesis can be tested in near future with the fast-evolving super-resolution imaging methods that are constantly pushing the boundaries of spatial and temporal resolutions of imaging and diffusion measurements on cell membrane.^97,98^

**Table 1:** Details of All Atom trajectories used to extract interaction energies

| System | Composition | # lipids | Temperature (K) | Duration (μs) |
| --- | --- | --- | --- | --- |
| DPPC/DOPC/CHOL | 0.37/0.36/0.27 | 360 | 298 | 9.43 |
| PSM/DOPC/CHOL | 0.43/0.38/0.19 | 452 | 295 | 4.54 |
| PSM/POPC/CHOL | 0.47/0.32/0.21 | 506 | 295 | 6.88 |

## Acknowledgement

The authors thank the D.E. Shaw Research (Anton Supercomputer) for making available the trajectories from Lyman’s group for our analysis. A.S. thanks the startup grant provided by the Ministry of Human Resource Development of India and the early career grant from the Department of Science and Technology of India. The authors thank Dr. Madhusmita Tripathy for useful insights into lattice models.

## Supporting Information Available

The following files are available free of charge.

- Filename: brief description
- Filename: brief description

This material is available free of charge via the Internet at http://pubs.acs.org/.

## Graphical TOC Entry

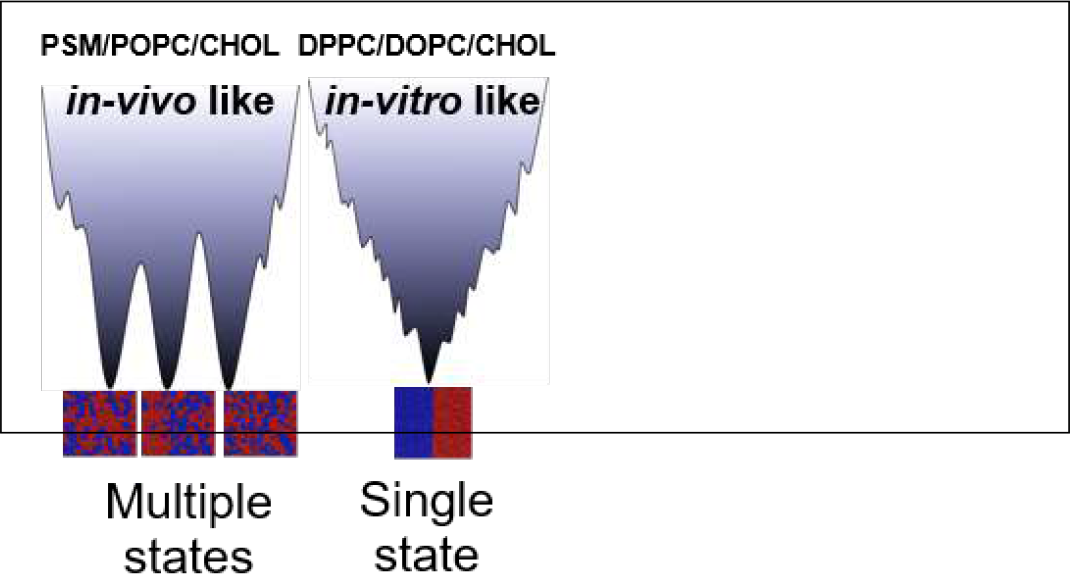

## Supporting Information

### 1 Description of systems studied

The three All Atom (AA) *L_o_/L_d_* phase co-existing systems studied and their trajectory details are given in Table 1 in the main text. These systems were simulated on Anton and the trajectory was made available by Dr. Edward Lyman and Anton.

DPPC/DOPC/CHOL [1] – An initial phase separated starting configuration of DPPC/DOPC/CHOL system was created by embedding an equilibrated circular *L_o_* patch in a *L_d_* system. When the system is equilibrated, the *L_o_* lipids in the circular patch attain a hexagonal packing with cholesterol and *L_d_* lipids forming the sea. The *L_o_* lipids region is manifested as the smaller cluster of *χ*^2^ values shown in Figure S2.

PSM/DOPC/CHOL and PSM/POPC/CHOL [2]- Initial configuration for these systems was generated using CHARMM-GUI. These systems phase separate in the time frame of their evolution into *L_o_* and *L_d_* phases but have very complex lateral geometries as seen in Figure S2 top panel where lipids are colored based on their chemical identity.

The systems were simulated using CHARMM36 force field [3] in NPT ensemble. Since all the three AA systems have symmetric membrane, the analysis was performed on one of the monolayers for each system.

### 2 Non-affine measurements in ternary lipid systems

Measure of non-affiness in the topological rearrangements (*χ*^2^) of lipid sites of an evolving system is a quantification of degree of non-uniform strain propagation in its local environment. In our previous work we have shown that *χ*^2^ can be used as a measure of “orderliness” in lipid systems [4]. The non-affine parameter characterizes the local environment of the lipid species. The formalism used to measure *χ*^2^ is similar to that used in the original work by Falk and Langer in 1998 [5]:

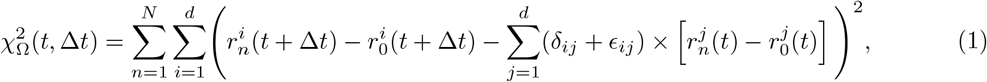

where the indices *i* and *j* run through the spatial coordinates for dimension *d* and *n* runs over the *N* lipids in the neighbourhood Ω, defined within a cutoff distance around the reference lipid *n* = 0. We define Ω to be the distance till second shell in the radial distribution of P atom sites – 14 *Å. δ_ij_* is the Kroneker delta function. *ϵ_ij_* is the strain associated with the maximum possible affine part of the deformation and thus, minimizes 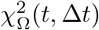. *ϵ_ij_* and 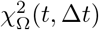 can be calculated following the method given in the original reference. Δ*t* is taken to be 240 ps in all the three systems as this was the dump frequencies of these trajectories.

Lipid systems in a single phase have a unimodal distribution of *χ*^2^ values, with systems in *L_o_* phases having lower *χ*^2^ values in comparison to systems in *L_d_* phase. Systems with liquid-liquid phase coexistence has a bimodal distribution of *χ*^2^ values as seen in Figure S3. Further *χ*^2^ values capture nuances of complex geometrical structures formed in phase separated lipid systems and can also be used to identify boundaries between phases. This sensitive nature of *χ*^2^ values and its ability to identify phases without being biased by the chemical identity of lipids makes it a ideal parameter to monitor complex geometries in lateral organization in lipid bilayers.

Figure S2 shows a comparison of contour map of P atom sites coloured based on their *χ*^2^ values and the corresponding configuration of lipids colour coded based on the lipid identity clearly shows that most P atoms with low *χ*^2^ values are mapped to the saturated lipids (rendered in red) and while most P atoms with higher *χ*^2^ values mapped to the unsaturated lipids (rendered in blue). Saturated lipids which are more dispersed among the unsaturated lipids have higher *χ*^2^ values. Hence these lipids are classified into *L_d_* phase lipids. DPPC/DOPC/CHOL system has a small tightly clustered *L_o_* lipids in the sea of *L_d_* lipids. Phase separation is most apparent in PSM/DOPC/CHOL system in comparison to a more heterogeneous spatial distribution of *χ*^2^ values in PSM/POPC/CHOL system.

### 3 Comparison of *χ*^2^ against *S_cd_* and *ϕ*_6_ order parameters

To highlight the advantage of using *χ*^2^ over existing characterization parameters to identify molecular-level phase co-existence in the CG and AA systems, we calculate hexatic order parameter (*ϕ*_6_) (for CG and AA systems Figure S4a) and deuterium order parameter (*S_cd_*) (for AA systems Figure S4b) and compare it to phases identified by *χ*^2^ characterization.

*ϕ*_6_ is calculated for the six nearest neighbours of the central atom. Its given as [6]:

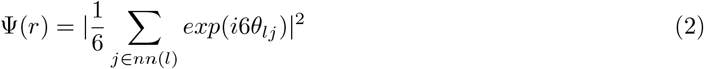

here *θ_ιj_* is the angle between reference axis and a vector connecting particle *l* to particle *j* and the summation *j* ∈ *nn*(1) is over the six nearest neighbours of particle *l*. 〈Ψ〉 is unity for a perfectly hexagon packing and zero when hexagonal packing is completely absent.

*S_cd_*, measure the flaccidity of the lipid alkyl chain tails, is the second order Legendre polynomial *P*_2_(*cosθ*):

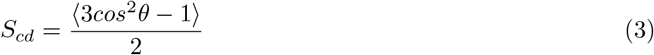

here *θ* is the angle between C-H bond vector and the bilayer normal and angular brackets denote ensemble average. S_cd_ values range between 1 (highly ordered) to −0.5 (disordered). Calculation of S_cd_ requires the knowledge of position of all the C-H vectors of the alkyl chain while calculating *χ*^2^ requires the position of any site from each lipid monitored with time. Hence *χ*^2^ has a dynamic nature hidden in its formalism.

Figure S4a shows the comparison between *φ_6_* and *χ*^2^ to identify *L_o_* and *L_d_* phases in DUPC/DPPC/CHOL CG system. The image on the left shows domains as identified by the chemical identity of lipids. While lipids corresponding to different phases are easily identified using *χ*^2^ values, the phases are less apparent using *φ*_6_ order parameter. Figure S4b shows a comparison of *φ*_6_, *S_cd_* and *χ*^2^ to define molecular-level phases in the PSM/DOPC/CHOL AA system. While *φ*_6_ is a good order parameter to distinguish packing in gel and fluid phase, as seen in Figure S4b, it fails to characterize *L_o_* and *L_d_* phases in fluid phase coexisting systems. S_cd_ and *χ*^2^ values are anti-correlated [4]. While both these quantitates can be used to distinguish *L_o_* and *L_d_* phases, as discussed above, calculating *χ*^2^ is computationally less expensive and also has a hidden dynamical nature. Hence we use *χ*^2^ values to mark *L_o_* and *L_d_* lipids and use them as lattice site parameters over the other existing methods to characterize *L_o_* and *L_d_* phases in the simulation and experiment literature.

### 4 Curating interactions from AA trajectories and mapping it to higher dimensions

To understand the driving force behind these complex organization patterns, lipid interaction energies are calculated by re-running long trajectories of atomic resolution systems simulated using CHARMM36 force field for lipids [3] using the re-run utility on Gromacs 5.1.2 [7]. Interaction energies including van der Waals and electrostatic contributions, for each lipid is calculated within a cut-off of 14*Å*, involving lipids present within the second nearest neighbour distance of the reference lipid. Figure S1b is a pictorial representation of the procedure followed to extract interaction energies per lipid. Circles demote the regions within which van der waal and coulombic interaction of the central lipid is calculated. The extracted energies are averaged over last 80 ns of the trajectory to ensure the values thus obtained were for systems at thermal equilibrium. The interaction energy thus obtained for each lipid is plotted as a function of its *χ*^2^ values in figure S5.

### 5 Lattice evolution

Regular simulated annealing (SA) algorithm is used for lattice evolution of these systems with continuous energy landscapes [8]. SA is a useful optimization method for these systems with continuous energy landscapes as it ensures extensive configurational search for all possible arrangements of lattice sites and minimization to a global minima configuration. The algorithm implemented in finding the global minima is as follows. Initial starting lattice configuration contains *χ*^2^ values randomly arranged in the lattice sites. Two arbitrary lattice points are picked and their positions are exchanged. The energy change incurred in this lattice rearrangement is calculated and the Metropolis Monte Carlo criteria is employed in accepting or rejecting the particular lattice swap move. Swapping *χ*^2^ values ensures that its total probability distribution is conserved during the lattice evolution. Initial MC sampling is performed at high temperatures which increases the probability of swaps being accepted. This ensures proper sampling of the configuration space. The temperature is then slowly decreased and MC sampling is performed at each temperature until local minima is attained at that temperature. At lower temperatures the number of bad moves being accepted dramatically decreases and the system is locked in its minima. The heating and cooling cycles are repeated in order to ensure that the system has indeed attained a global minima configuration. As an additional check, to rule out the effect of starting configuration on the final *χ*^2^ organization patterns, parallel SA simulations are run from many different initial configurations, ranging from a completely phase separated configuration to random arrangement of *χ*^2^ values. The final lattice organization obtained from the set of parallel runs are compared.

### 6 PSM/DOPC/CHOL lattice configurations

Figure S6 shows the final lattice configuration for the PSM/DOPC/CHOL system from different initial configurations. The final configurations are similar to the phase separated *L_o_/L_d_* organization in the AA system.

### 7 Effect of changing the ratio of *L_o_*:*L_d_* lipids in lateral organization patterns

Population distribution of *L_o_* and *L_d_* lipids with varying *χ*^2^ values for DPPC/DOPC/CHOL and PSM/POPC/CHOL system are shown in figure S7 and figure S9 respectively. Figure S8 shows the invariance in lattice organization in DPPC/DOPC/CHOL system with different ratio of *L_o_*:*L_d_* lipids.

**Figure S1a:**
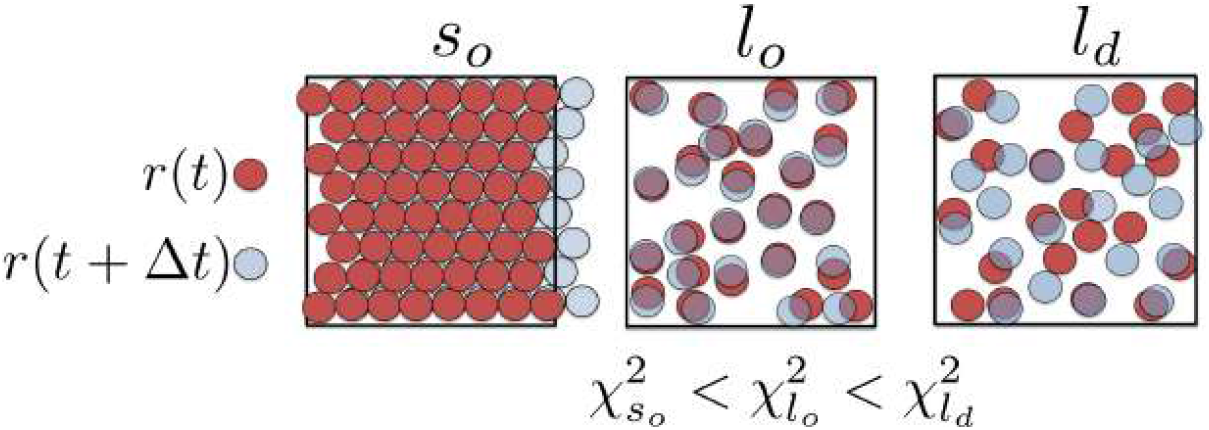
Schematic showing how *χ*^2^ measures topological rearrangements in atom sites to distinguish between phases in lipid systems. Red and blue circles represent position of (any) lipid atom site at position *t* and *t* + Δ*t*

**Figure S1b:**
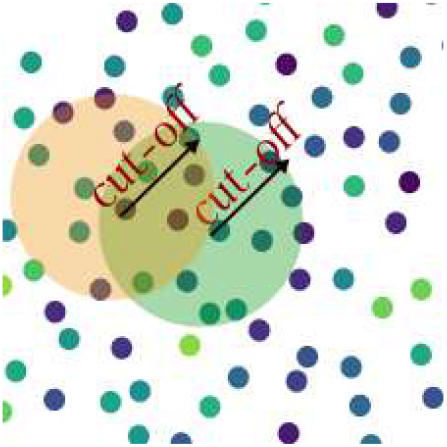
Pictorial representation of calculating interactions for AA trajectories. Circles demote the regions within which van der waal and coulombic interaction of the central lipid is calculated.

**Figure S2:**
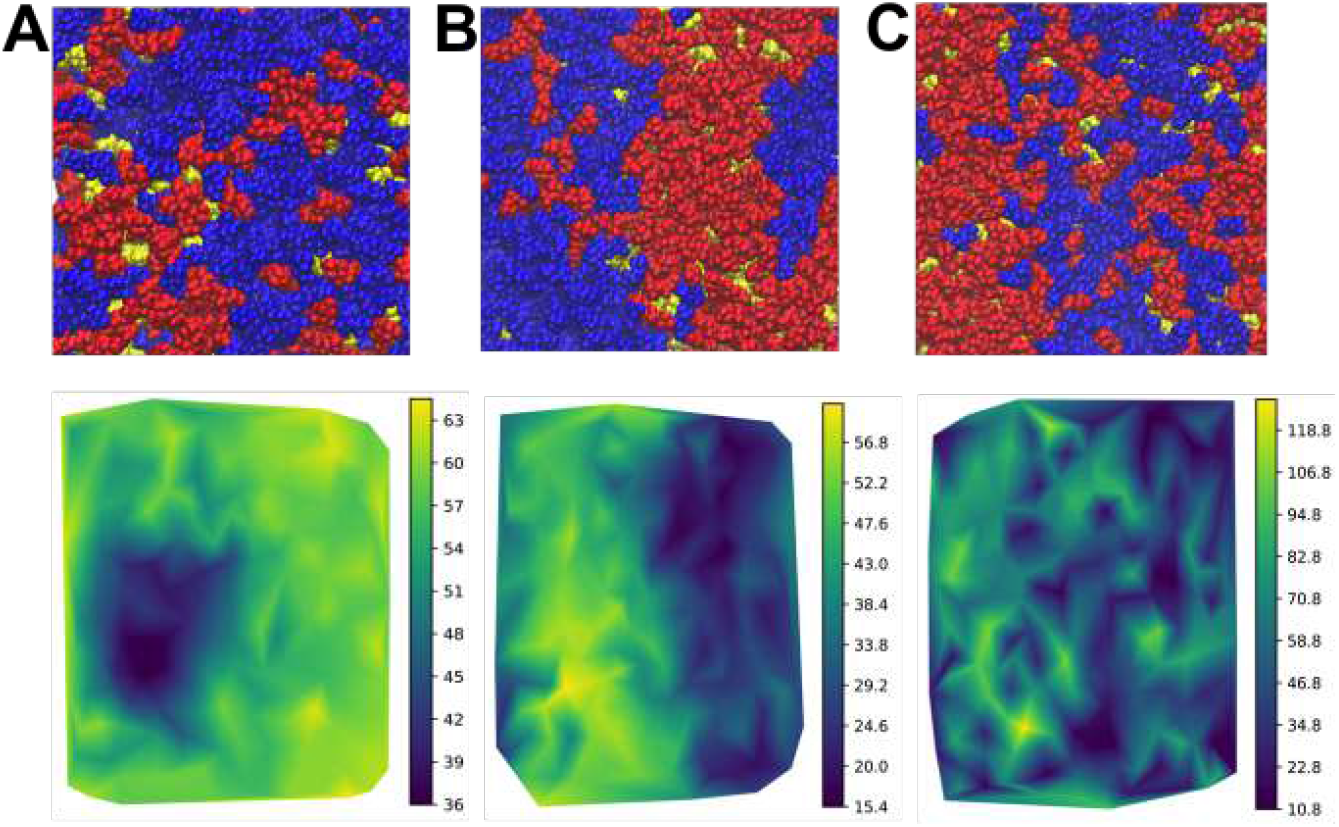
final configuration of (A) DPPC/DOPC/CHOL, (B) PSM/DOPC/CHOL and (C) PSM/POPC/CHOL systems with red colour for lipids with saturated alkyl chain tails (PSM/DPPC), blue for lipids with unsaturated alkyl chain tails and yellow for cholesterol. Image below shows contour map of P atom sites colored based on *χ*^2^ for the respective systems.

**Figure S3:**
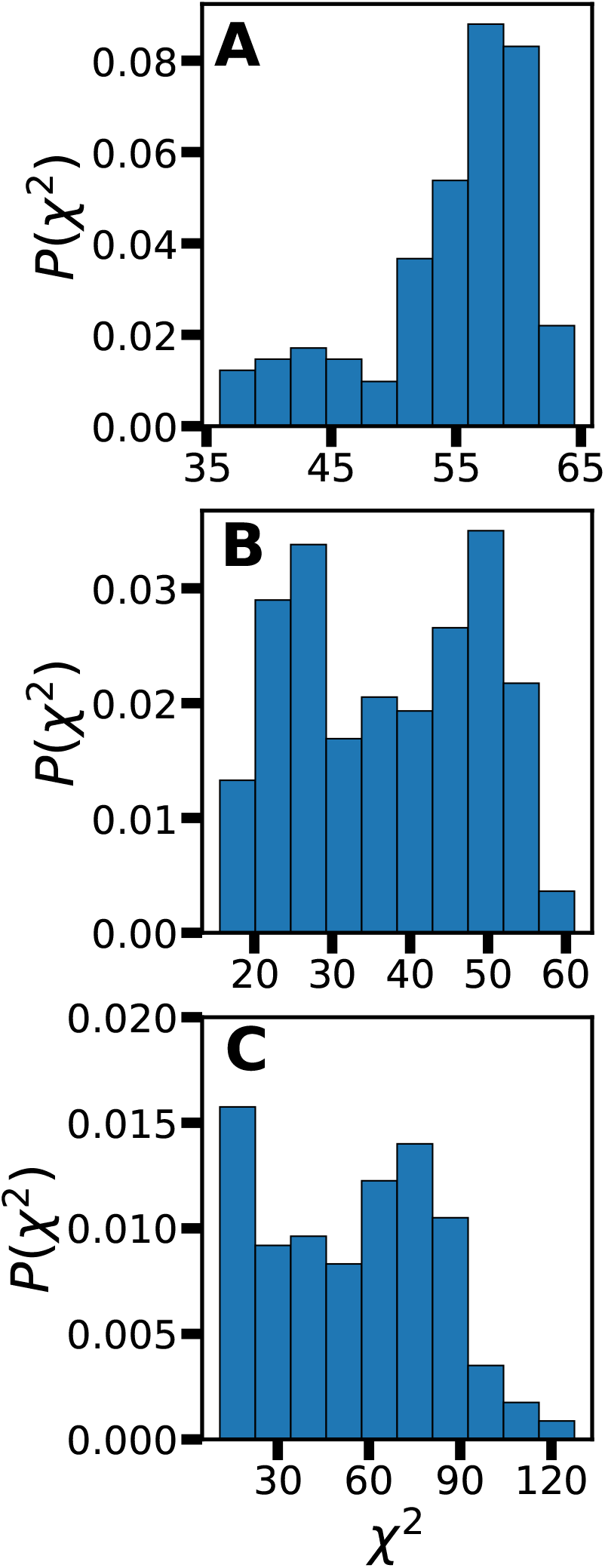
Bimodal probability distribution of *χ*^2^ values for phase separated (A) DPPC/DOPC/CHOL, (B) PSM/DOPC/CHOL and (C) PSM/POPC/CHOL systems.

**Figure S4a:**
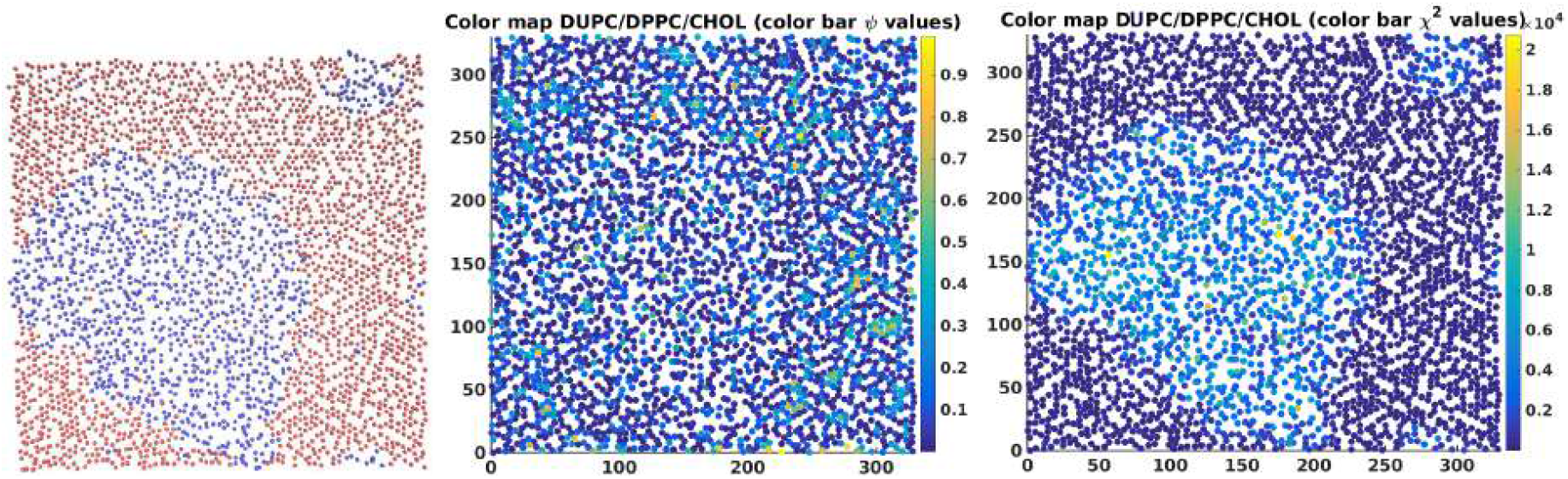
Comparison of hexatic order parameter (middle) and *χ*^2^ values (right) for coarse grain (CG) DPPC/DUPC/CHOL system. Image on the (left) shows mid-C atoms of the alkyl chains color coded based on their chemical identity (DPPC-red and DUPC-blue).

**Figure S4b:**
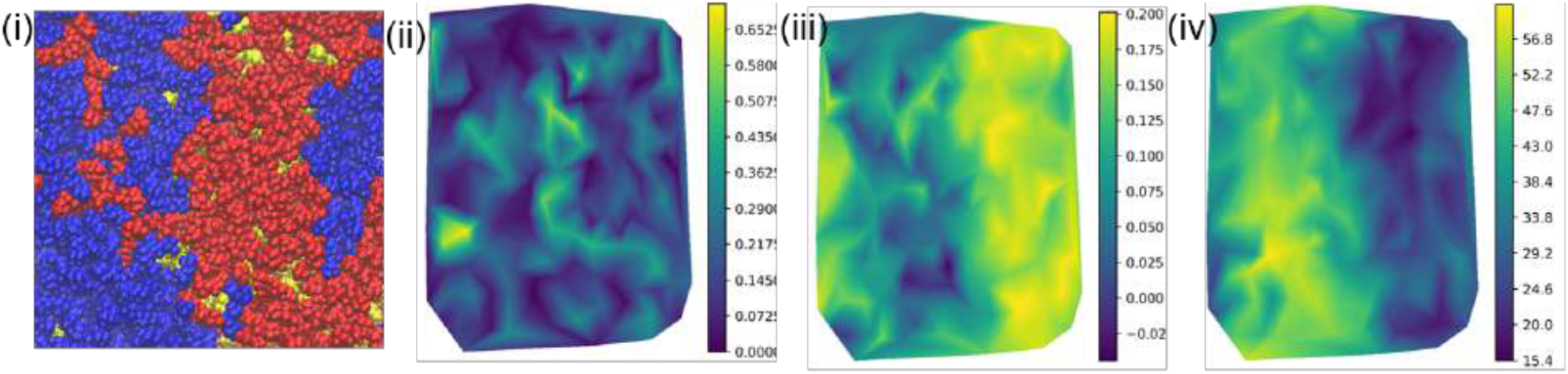
Contour plot of lipid P atom sites coloured based on the (ii) Hexatic order parameter (*ϕ*_6_), (iii) Tail order parameter (*S_cd_*) and non-affine parameter (*χ*^2^) for PSM/DOPC/CHOL system. (i) shows the lipid sites colored based on their chemical identity (PSM-red DOPC-blue and cholesterol – yellow).

**Figure S5:**
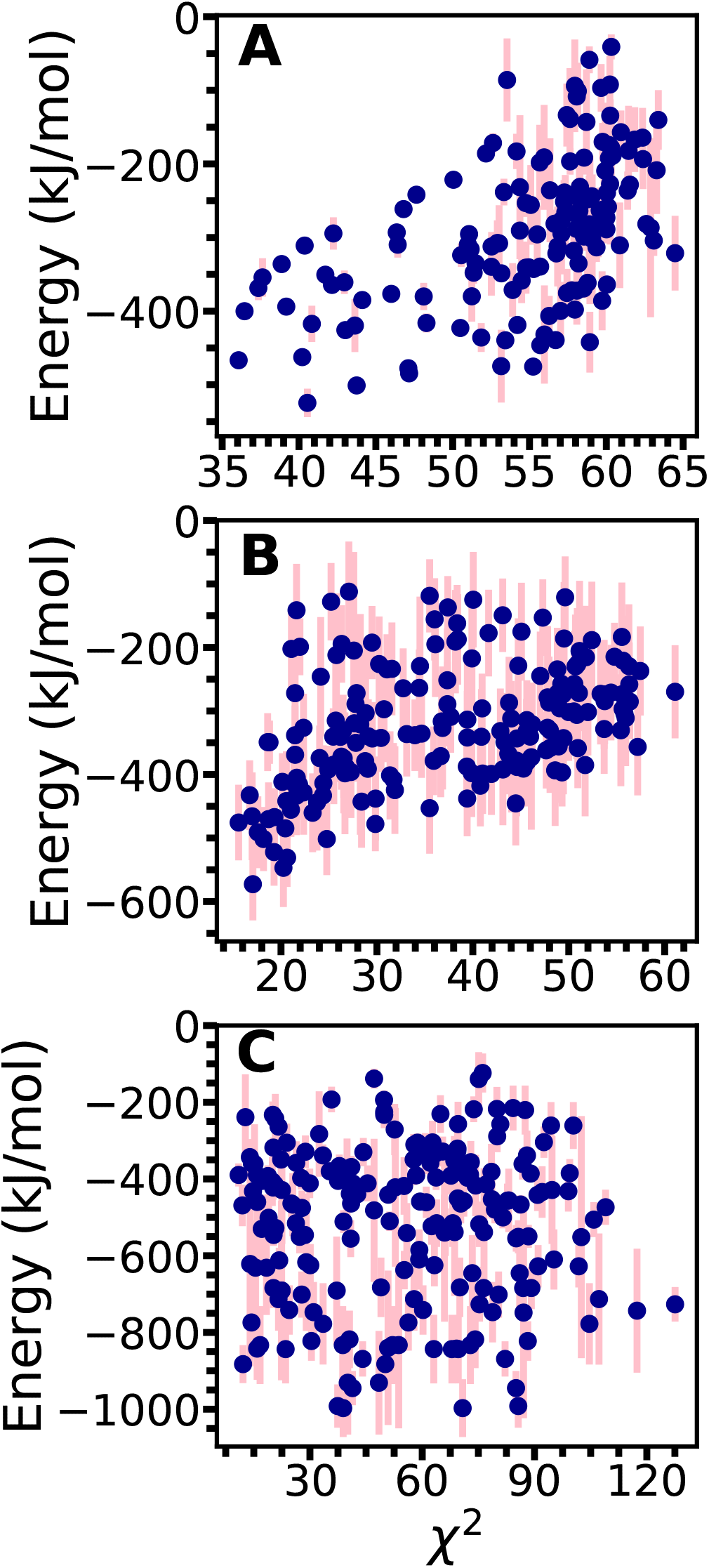
Interaction energy plotted as a function of *χ*^2^ values for (A) DPPC/DOPC/CHOL, (B) PSM/DOPC/CHOL and (C) PSM/POPC/CHOL. The blue points and pink lines indicate average interaction energy values and its standard deviation calculated is the last 80 ns of the evolution.

**Figure S6:**
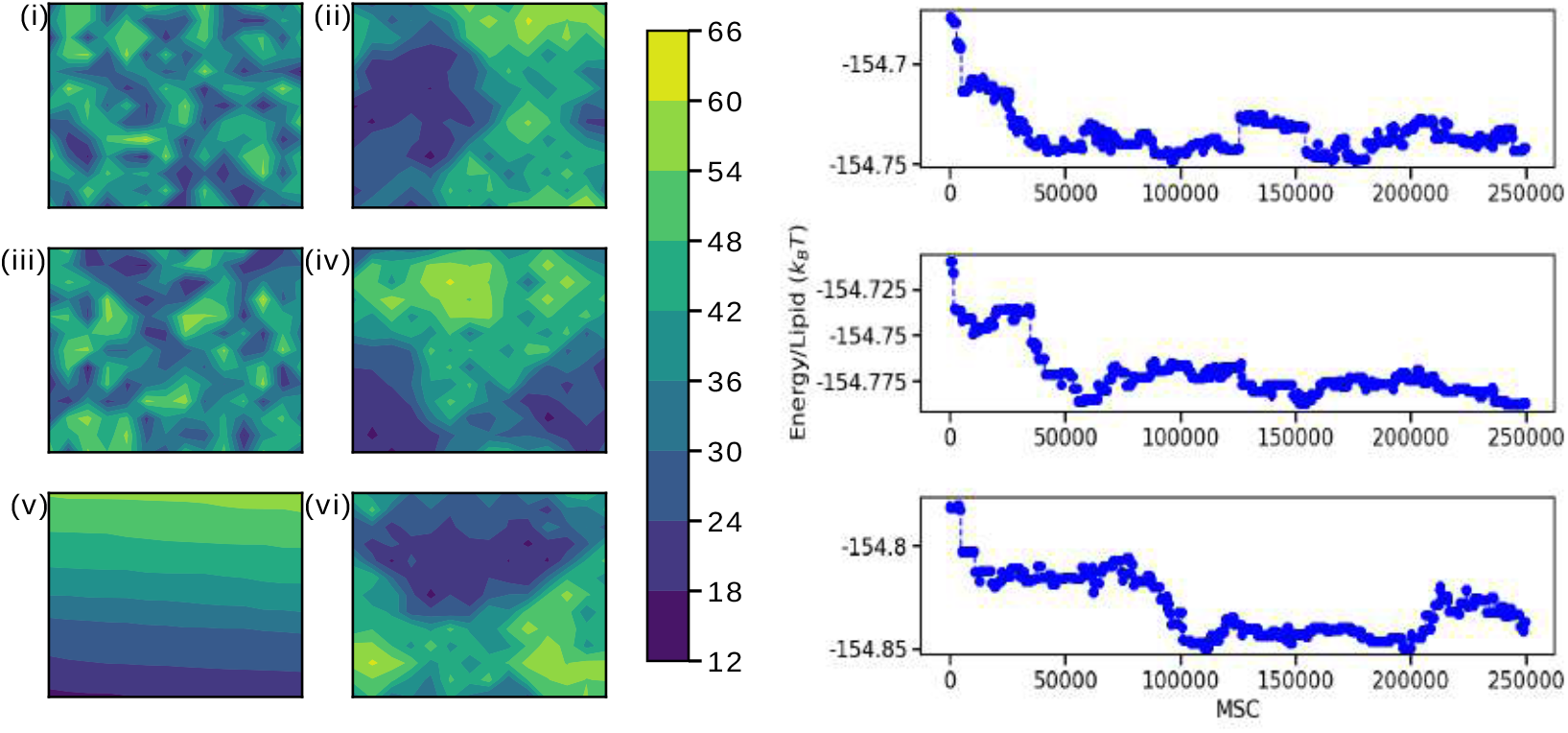
Initial (i,iii,v) *χ*^2^ organization and final (ii,iv,vi) *χ*^2^ organization for PSM/DOPC/CHOL mixture. (i)-(ii) and (iii)-(iv) represent lattice evolutions starting from random lattice arrangements. (v)-(vi) represent lattice evolution from a phase separated arrangement of *χ*^2^ values. The sites have been color coded according to the *χ*^2^ values. The Energy/lipid values for the last 2.5 * 10^5^ MCS evolution corresponding to evolutions is shown in the right. Similar energies show that these states are degenerate.

**Figure S7:**
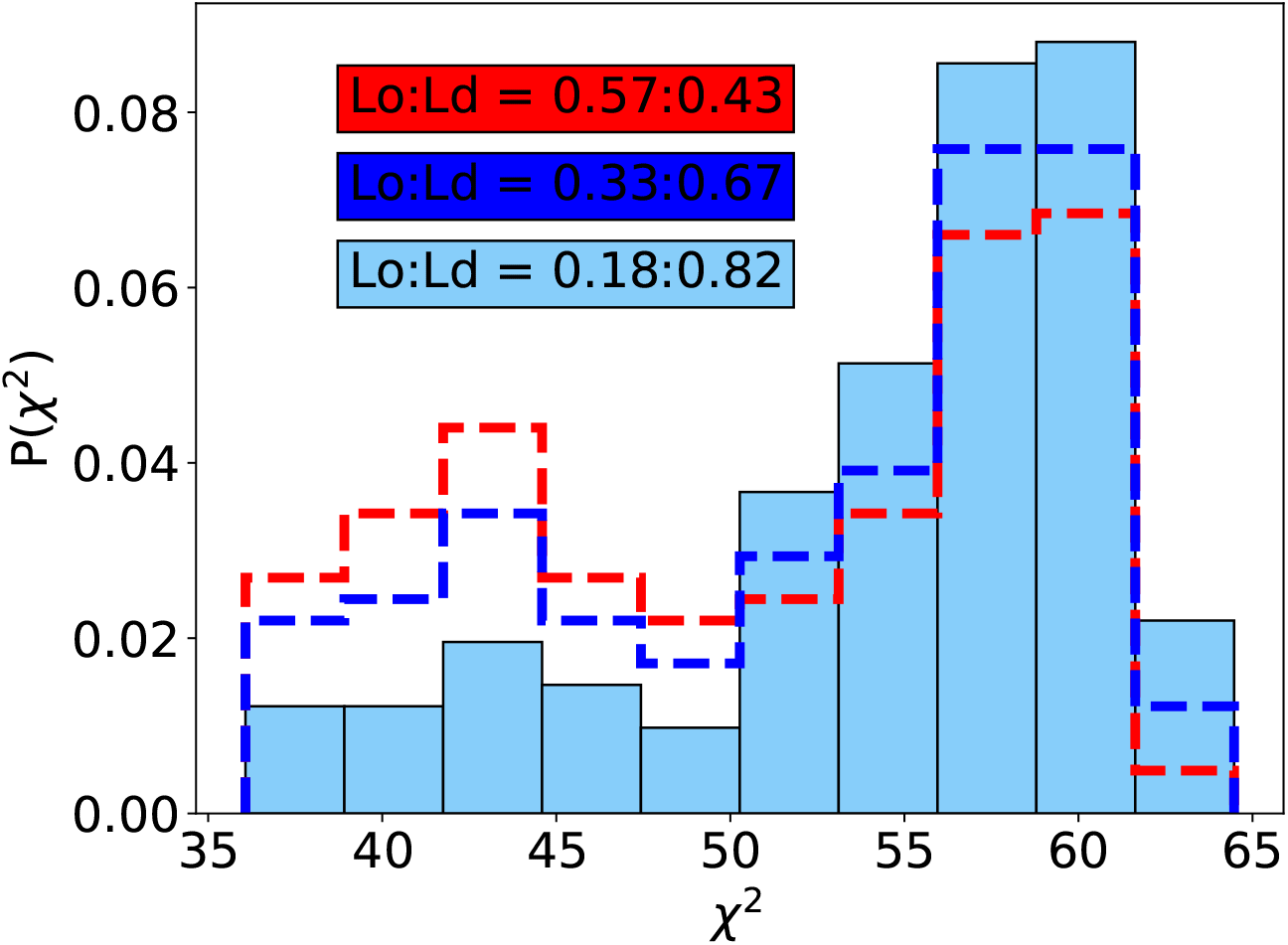
Histogram of *χ*^2^ values with varying ratios of *L_o_*:*L_d_* lipids for DPPC/DOPC/CHOL system.

**Figure S8:**
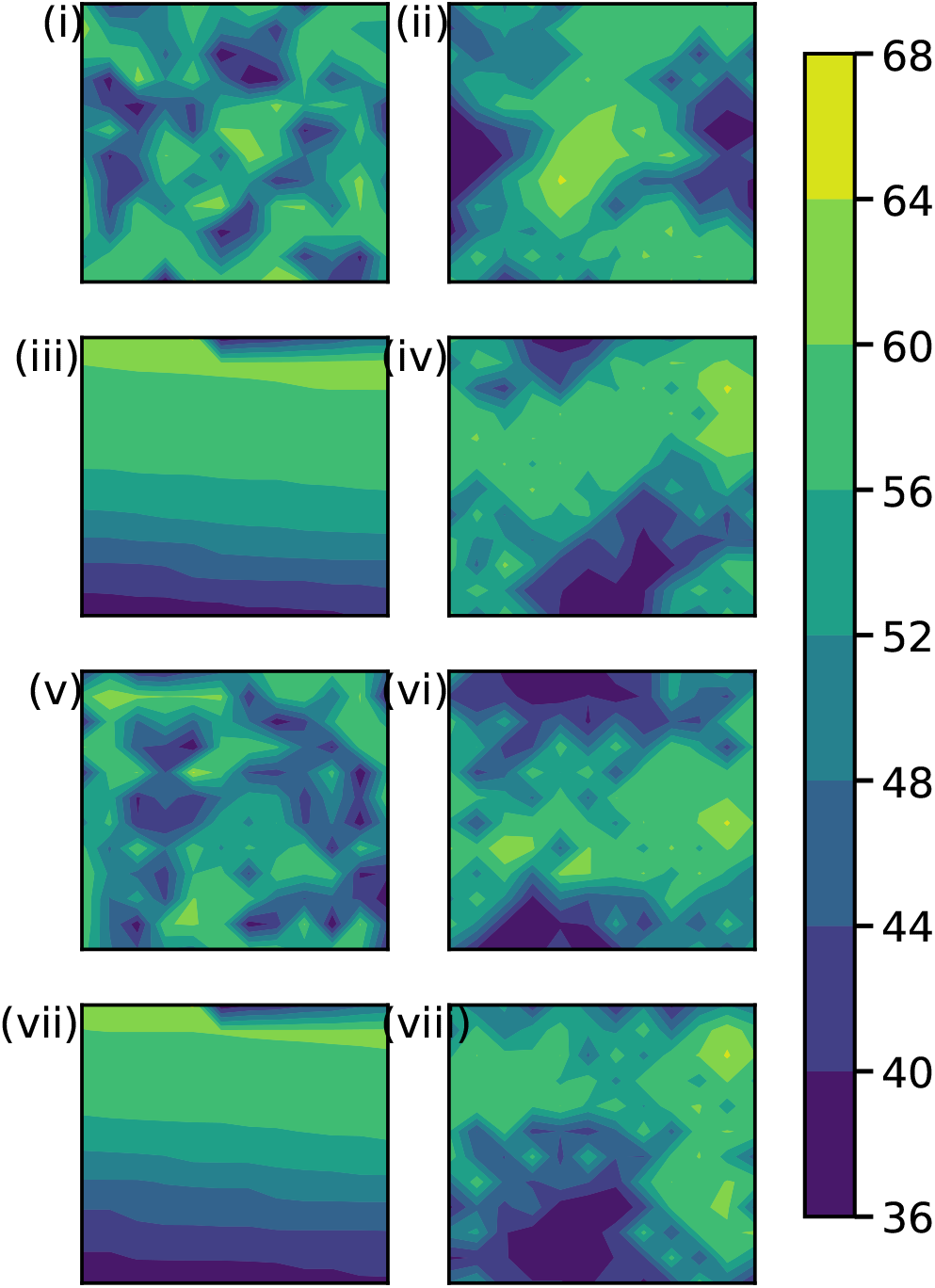
Initial (i,iii,v,vii) *χ*^2^ organization and final (ii,iv,vi,viii) *χ*^2^ organization for DPPC/DOPC/CHOL mixture. (i)-(iv) represent lattice evolutions starting from random lattice arrangements (i-ii) and phase separated lattice arrangement (iii-iv) for *L_o_:L_o_* ratio of 0.33:0.67. (v)-(viii) represent lattice evolutions starting from random lattice arrangements (v-vi) and phase separated lattice arrangement (vii-viii) for *L_o_:L_o_* ratio of 0.57:0.43. The sites have been color coded according to the *χ*^2^ values.

**Figure S9:**
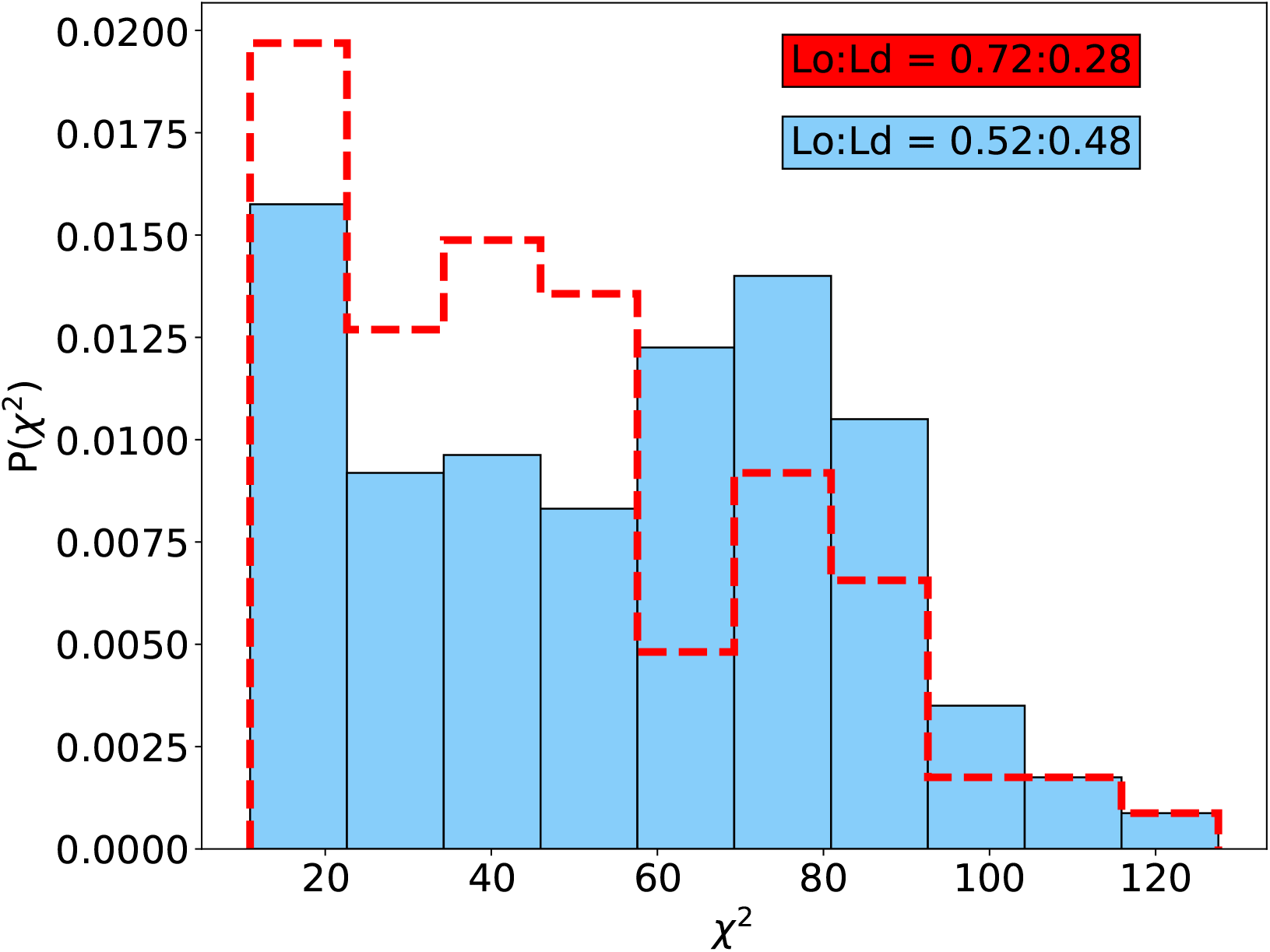
Histogram of *χ*^2^ values with varying ratios of *L_o_*:*L_d_* lipids for PSM/POPC/CHOL system.

